# Behavioral test batteries induce transient, domain-specific effects while preserving global phenotypic structure in zebrafish

**DOI:** 10.64898/2026.08.17.745208

**Authors:** Barbara D. Fontana, Camilla W. Pretzel, Mariana M. Schmitz, Mariana L. Müller, Angela E. Uchoa, Eduarda T. Saccol, Cássio M. Resmim, Denis B. Rosemberg

**Affiliations:** Laboratory of Experimental Neuropsychobiology, Department of Biochemistry and Molecular Biology, Natural and Exact Sciences Center, Federal University of Santa Maria, Santa Maria, Brazil; Graduate Program in Biological Sciences: Toxicological Biochemistry, Federal University of Santa Maria, Santa Maria, Brazil; The International Zebrafish Neuroscience Research Consortium (ZNRC), Slidell, USA

**Keywords:** anxiety, novel tank test, phenotyping, sequential testing, social behavior

## Abstract

Behavioral test batteries are increasingly used to characterize multiple functional domains in zebrafish, yet the potential impact of test sequence on behavioral outcomes remains poorly defined. Here, we systematically evaluated whether test order influences behavioral responses in a three-assay battery comprising the novel tank test (NTT), mirror-induced aggression (MIA), and social preference (SP) test. Adult zebrafish (*Danio rerio*) were exposed to all possible permutations of the three assays in a fully counterbalanced design, allowing assessment of order effects across locomotor, anxiety-like, aggression-related, and social behaviors. Test order produced modest and parameter-specific effects, primarily affecting locomotor activity in the NTT and social proximity in the SP assay. Time-course analysis revealed within-test behavioral dynamics, with limited evidence that test order modulates early adaptation or late engagement with the testing environment but does not alter overall temporal response profiles. Sex-dependent effects were assay-specific and most pronounced in the NTT, with no consistent sex differences observed in MIA or SP. To evaluate the global structure of behavioral variation, Principal Component Analysis (PCA) was performed across assays. Despite localized effects of test order, no clear multivariate separation between test sequences was observed, indicating that sequential testing does not produce distinct baseline phenotypes. Together, these findings support the robustness and reproducibility of multidomain behavioral batteries while highlighting the importance of standardized test-order reporting to improve cross-study comparability.

## Main

Behavioral phenotyping in adult zebrafish (*Danio rerio*) has expanded considerably, with multiple validated paradigms targeting distinct neurobehavioral domains, including anxiety-like behavior^1–3^, social interaction^4–6^, aggression^7,8^, and cognition^9–11^. This species has become a widely used vertebrate model in translational neuroscience due to the conserved neurobiology, genetic tractability, and suitability for medium– to high-throughput behavioral screening^12^. Zebrafish display a complex behavioral repertoire^13,14^ spanning numerous measurable endpoints across functional domains. Accordingly, various assays have been developed to quantify specific behavioral responses, including the novel tank test (NTT)^1,3^, social preference (SP)^4,15^, and mirror-induced aggression (MIA)^8,16^, which assess complementary constructs such as novelty-induced anxiety, affiliative motivation, and aggression-related responses.

Because neurobehavioral phenotypes are inherently multidimensional, several studies combine multiple assays within the same experimental cohort to obtain broader behavioral characterization. This multidomain approach, often referred to as behavioral test batteries, parallels established strategies in rodent behavioral neuroscience, where sequential testing is used to capture and correlate complex phenotypes^17,18^. Recent analyses of zebrafish behavioral literature indicate that multi-assay experimental designs are increasingly common in adult zebrafish research, with approximately 79% of studies published in 2025 that incorporated the NTT also featuring additional behavioral assays, particularly anxiety-like, social, cognitive, and aggression-related paradigms^19^. These behavioral test batteries aim to provide broader multidomain phenotypic characterization within the same experimental cohort while improving experimental throughput and reducing cohort-related variability. In addition, their successful implementation may contribute to experimental refinement and reduction in animal use by allowing multiple behavioral domains to be assessed within the same individuals rather than requiring separate cohorts for each assay. However, sequential exposure to multiple behavioral tasks may introduce confounding effects, including stress carryover, habituation to novelty, and shifts in arousal or motivation that influence performance in subsequent assays^20,21^. Such order effects are well documented in rodents, where prior exposure to novelty– or stress-inducing tasks can significantly alter behavioral outcomes in later tests^20^. Thus, as multidomain behavioral batteries become increasingly common in zebrafish research, understanding the potential influence of test order is essential for ensuring experimental rigor, reproducibility, and cross-study comparability.

To address this question, we directly examined the impact of test order in a three-assay zebrafish behavioral battery composed of a multi-domain paradigm. Thus, naïve animals were individually exposed to NTT, SP, and MIA following a fully counterbalanced design that allows us to underscore whether prior exposure to one behavioral domain influences performance in subsequent assays and alters baseline behavioral responses **(Fig. 1)**. Finally, we evaluated potential sex-dependent effects, explored temporal patterns of behavior within assays, and used Principal Component Analysis (PCA) to examine the multivariate organization of behavioral phenotypes across the behavioral battery.

**Fig. 1.**
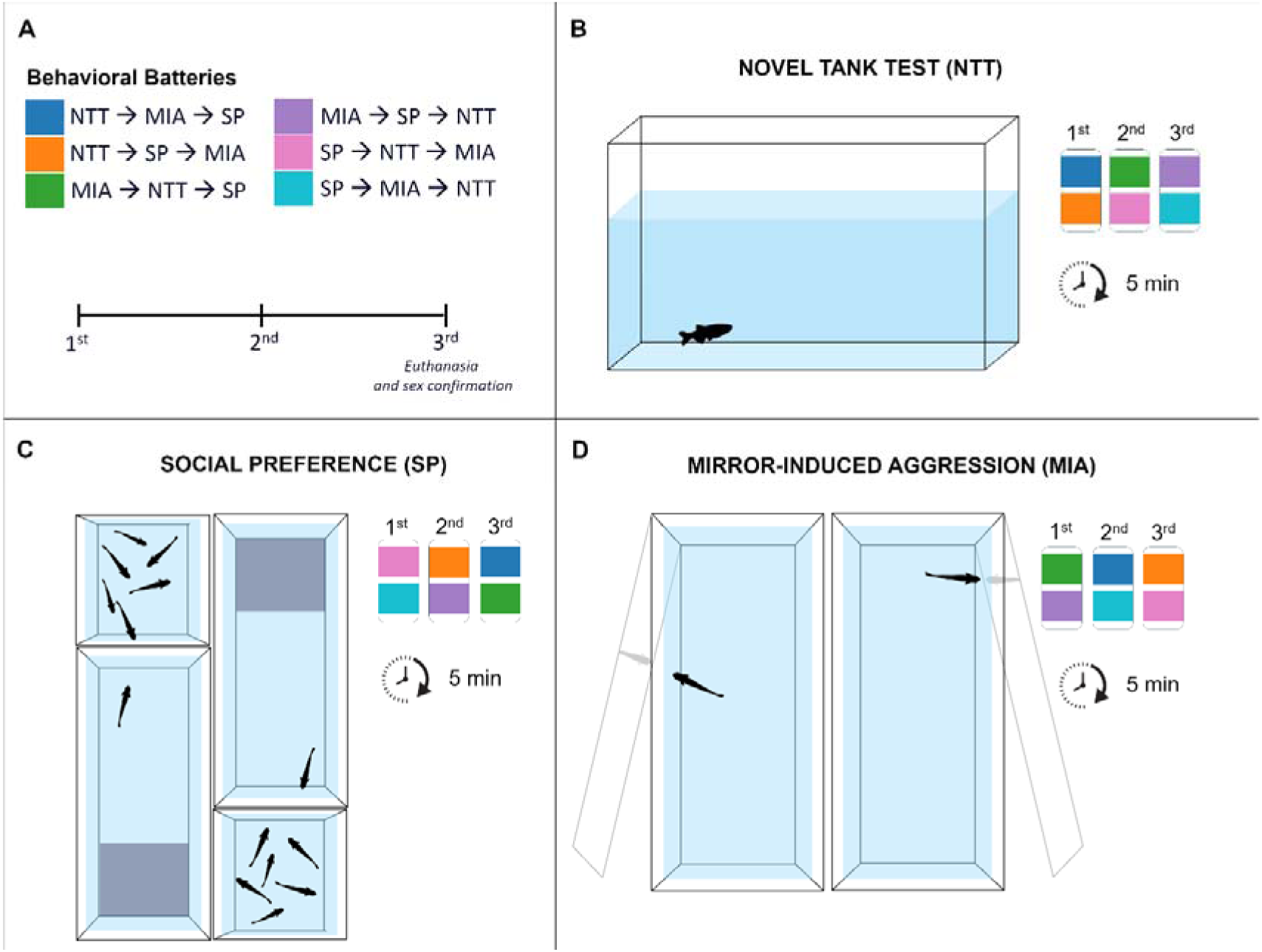
Experimental design and behavioral assay overview. **(A)** Schematic of the behavioral battery showing all test order permutations of the three assays: novel tank test (NTT), mirror-induced aggression (MIA), and social preference (SP). Animals were tested across three sequential positions (first, second, third), with colored blocks indicating the assay performed at each position. All possible assay orders were represented, in which colored blocks indicate the test position (first, second, third) for each experimental group. Following completion of the behavioral battery, animals underwent euthanasia and sex confirmation. **(B)** Novel tank test (NTT) setup. Individual fish were placed in a novel tank and behavior was recorded for 5 min. **(C)** Social preference (SP) test setup. Individual fish were tested in an experimental tank adjacent to a stimulus compartment containing six conspecifics. The tank was subdivided into virtual zones to quantify spatial preference relative to the social stimulus. **(D)** Mirror-induced aggression (MIA) test setup. Fish were exposed to a mirror stimulus positioned outside the tank, and behavior was recorded for 5 min to assess responses to the reflected image.

## Results

### Test order produces limited effects on behavioral endpoints

We first assessed whether test order influenced behavioral endpoints across the NTT, MIA, and SP assays using one-way ANOVA **(Fig. 2)**. In the NTT **(Fig. 2A)**, test order significantly affected locomotor activity. Both total distance traveled (*F*_(2,81)_ = 3.59, *p* = 0.032) and mean speed (*F*_(2,81)_ = 3.30, *p* = 0.042) differed across test positions. Post hoc comparisons indicated increased locomotor activity when the NTT was performed third compared to first (distance: *p* = 0.024; speed: *p* = 0.033). No significant effects of test order were observed for anxiety-related measures, including time spent in the top zone (*F*_(2,81)_ = 0.30, *p* = 0.743), mean distance from bottom (*F*_(2,81)_ = 0.63, *p* = 0.536), or boldness index (*F*_(2,81)_ = 0.20, *p* = 0.816).

**Fig. 2.**
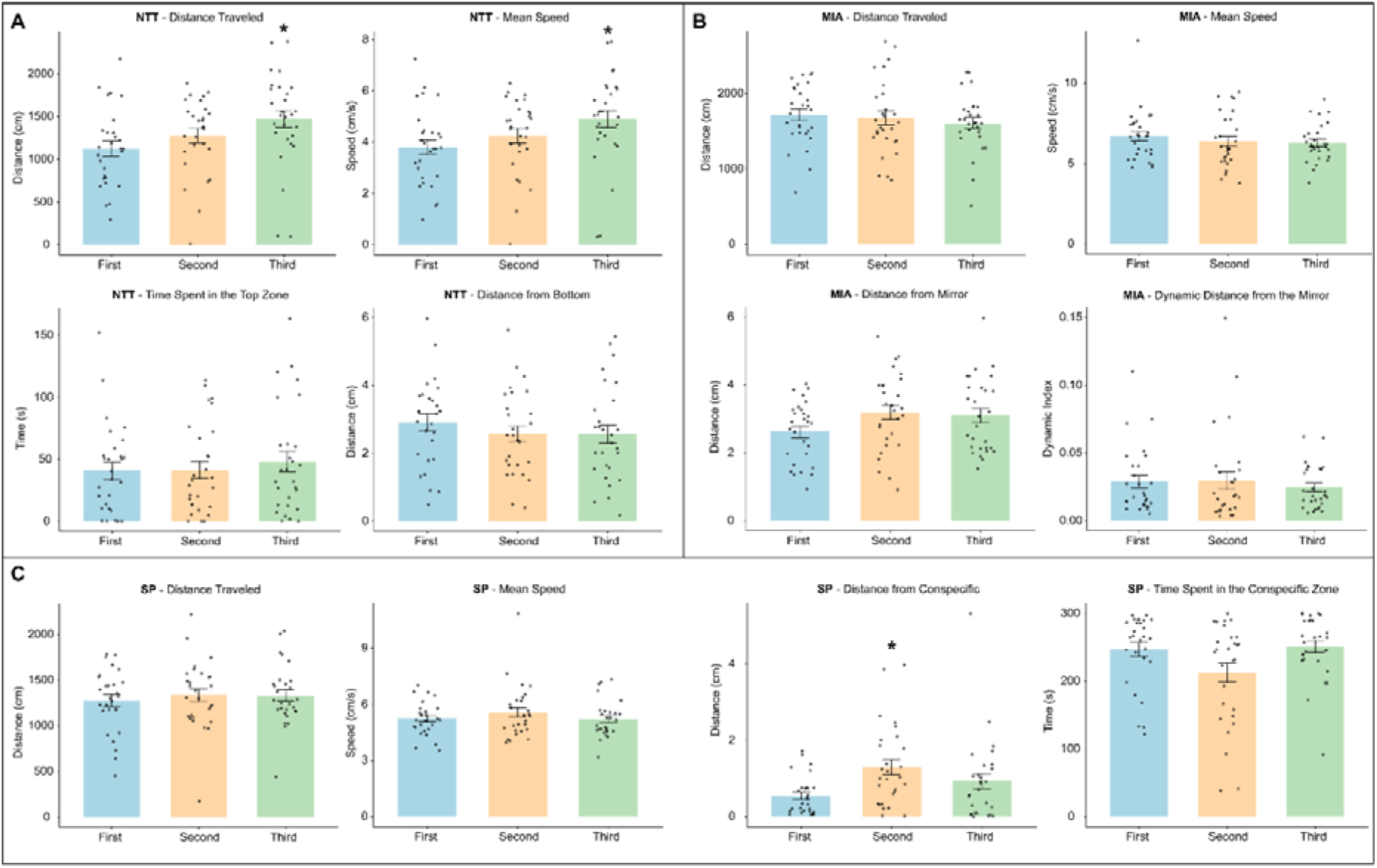
Behavioral endpoints across assays as a function of test order. **(A)** Novel tank test (NTT): total distance traveled (cm), mean speed (cm/s), time spent in the top zone (s), and mean distance from the bottom (cm) across test positions (first, second, third). **(B)** Mirror-induced aggression (MIA): total distance traveled (cm), mean speed (cm/s), mean distance from the mirror (cm), and dynamic aggression index. **(C)** Social preference (SP): total distance traveled (cm), mean speed (cm/s), mean distance to conspecifics (cm), and time spent in the conspecific zone (s). Data are presented as mean ± S.E.M., with individual data points shown. Data were analyzed using one-way ANOVA followed by Tukey’s post hoc test. Asterisks (*) indicate significant differences compared to first test order (*n* = 28; *p* <* 0.05).

In contrast, no significant effects of test order were detected in the MIA assay **(Fig. 2B)** across locomotor or aggression-related parameters (all *p* > 0.10). Similarly, in the SP assay, locomotor parameters were not affected by test order (all *p* > 0.40). However, a significant effect of test order was observed for distance to the social stimulus (*F*_(2,81)_ = 4.61, *p* = 0.013), with increased distance when the SP was performed second relative to first (*p* = 0.0097). No significant effects were observed for time spent near the stimulus (*F*_(2,81)_ = 3.42, *p* = 0.076) **(Fig. 2C)**. Analyses incorporating prior test exposure (“before” condition) revealed no consistent effects of prior exposure across behavioral parameters. While a small number of test order × prior exposure interactions reached significance, limited to NTT locomotor measures (distance traveled and mean speed) and MIA distance to the mirror (p < 0.05), all remaining parameters were unaffected and no coherent pattern emerged across assays **(Fig. S1)**. Overall, test order produced modest, parameter-specific effects without consistent effects across behavioral domains or a clear impact of prior test exposure.

**Fig. S1.**
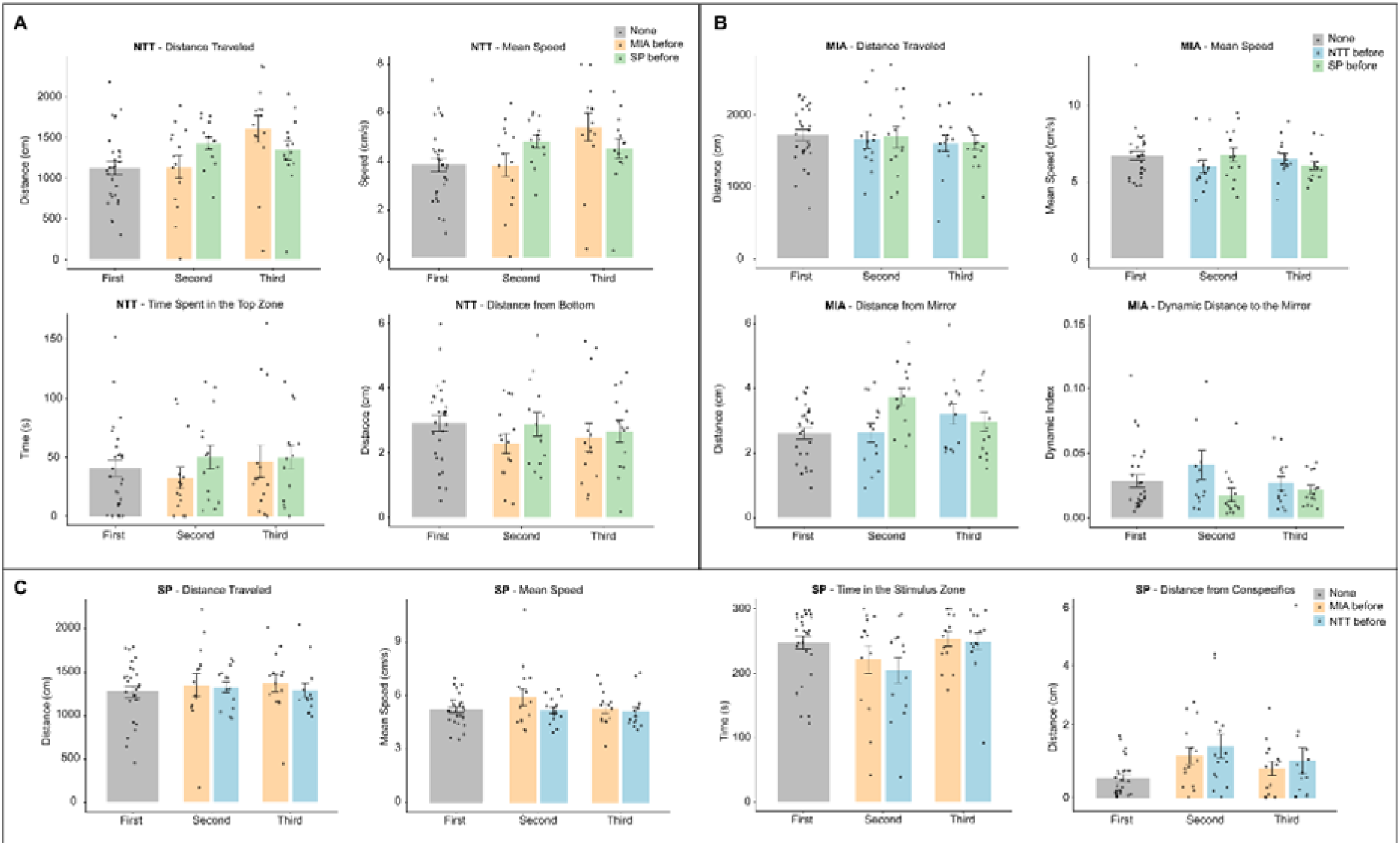
Effects of prior test exposure on behavioral endpoints across assays. **(A)** Novel tank test (NTT) behavioral parameters across test positions, including total distance traveled (cm), mean speed (cm/s), time spent in the top zone (s), and mean distance from the bottom (cm), stratified by prior exposure condition (none, MIA before, SP before). **(B)** Mirror-induced aggression (MIA) behavioral parameters across test positions, including total distance traveled (cm), mean speed (cm/s), mean distance to the mirror (cm), and dynamic aggression index, stratified by prior exposure condition (none, NTT before, SP before). **(C)** Social preference (SP) behavioral parameters across test positions (first, second, third), including total distance traveled (cm), mean speed (cm/s), time spent in the stimulus zone (s), and mean distance to conspecifics (cm), stratified by prior exposure condition (none, MIA before, NTT before). Data are presented as mean ± S.E.M., with individual data points shown, and were analyzed using two-way ANOVA with test position and prior exposure as factors, followed by Tukey’s post hoc test. Sample sizes ranged from *n* = 14–28 per group, with the “none” condition representing pooled observations from groups without prior exposure to the corresponding assay.

### Test order modulates behavioral adaptation without altering overall dynamics

To investigate potential carryover effects, behavioral dynamics were analyzed across the 5-min testing period using 30-s bins **(Fig. 3)**. Four representative parameters were selected to capture key behavioral domains: NTT distance traveled (locomotion), NTT mean distance from bottom (anxiety-like exploration), MIA distance to the mirror (aggression-related spatial behavior), and SP distance to conspecifics (social preference). All parameters exhibited significant time-dependent changes consistent with within-test behavioral adaptation (all *p* < 0.01). The main effect of test order was limited and parameter-specific. A significant effect of order was detected for NTT distance traveled (*F*_(2,81.07)_ = 3.12, *p* = 0.049) and SP distance to conspecifics (*F*_(2,81.39)_ = 3.91, *p* = 0.024), whereas no main effect of order was observed for NTT mean distance from bottom (*F*_(2,81.07)_ = 0.45, *p* = 0.636) or MIA distance to the mirror (*F*_(2,81.05)_ = 2.49, *p* = 0.089).

**Fig. 3.**
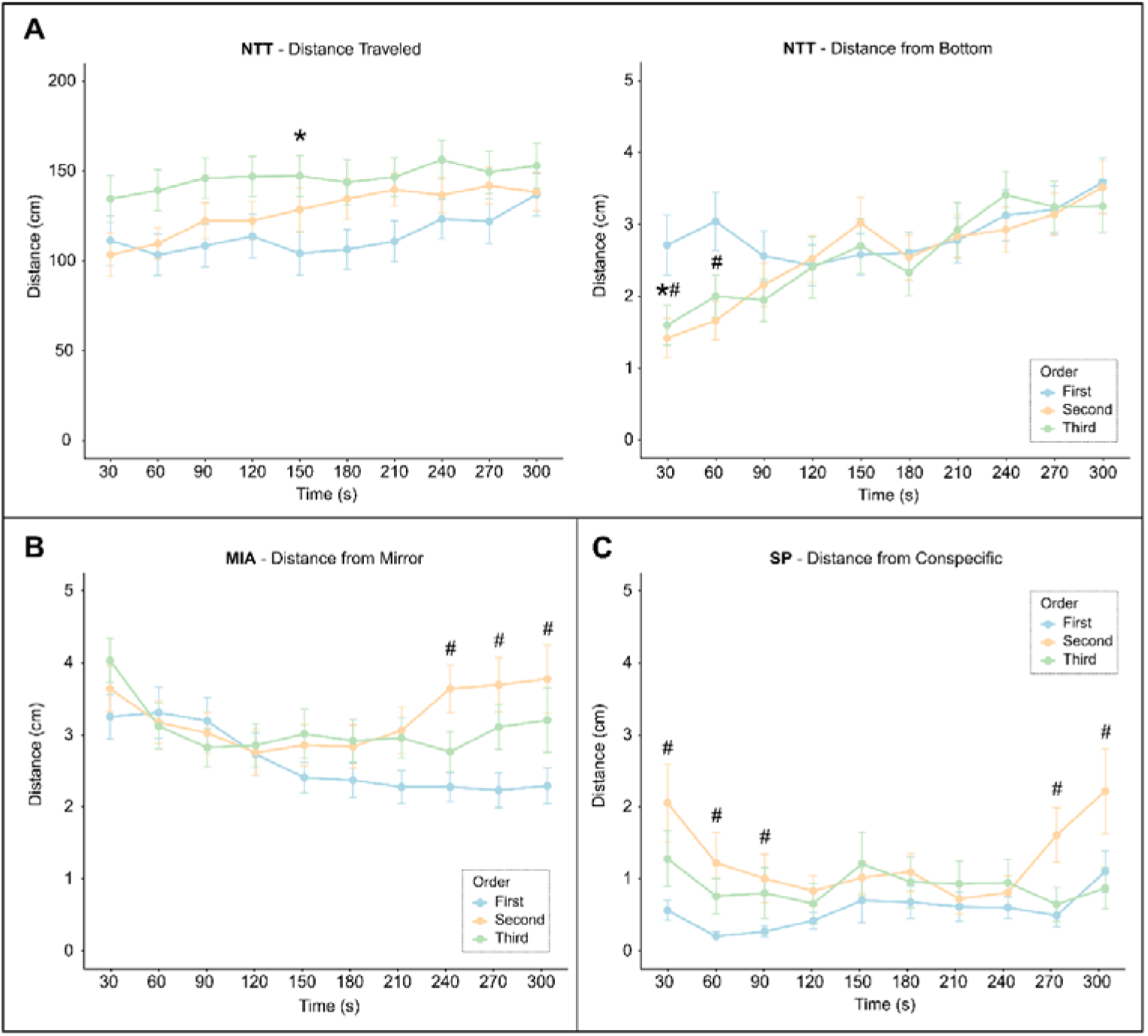
Temporal dynamics of behavior across assays as a function of test order. **(A)** Novel tank test (NTT) behavioral dynamics across the 5-min test, shown in 30-s bins, including total distance traveled (cm) and mean distance from the bottom (cm) across test positions (first, second, third). **(B)** Mirror-induced aggression (MIA) behavioral dynamics, represented by mean distance to the mirror (cm) across test positions. **(C)** Social preference (SP) behavioral dynamics, represented by mean distance to conspecifics (cm) across test positions. Data are presented as mean ± S.E.M. Data were analyzed using linear mixed-effects models with test order and time as fixed factors and subject as a random effect, followed by post hoc comparisons where appropriate. Asterisks (*) denote significant differences between first and third test positions, whereas hash symbols (#) denote significant differences between first and second test positions (*n* = 28; *p* <* 0.05).

Interactions between test order and time revealed that order-dependent effects were temporally restricted. Significant order × time interactions were observed for NTT mean distance from bottom (*F*_(18,721.21)_ = 2.09, *p* = 0.005) and MIA distance to the mirror (*F*_(18,721.26)_ = 2.16, *p* = 0.0036), while no interaction was detected for NTT distance traveled (*F*_(18,721.16)_ = 0.91, *p* = 0.565) or SP distance to conspecifics (*F*_(18,721.73)_ = 1.33, *p* = 0.161). Post hoc comparisons indicated that these interaction effects were primarily confined to early phases of the assays. In the NTT, fish tested first exhibited greater distance from the bottom at 30s and 60s compared to those tested second (30 s: *p* = 0.017; 60 s: *p* = 0.0096), and at 30s compared to those tested third (*p* = 0.047), indicating reduced bottom-dwelling during initial exposure. Similarly, in the SP assay, fish tested first in the assay battery displayed reduced proximity to conspecifics at early time points relative to those tested second (30 s: *p* = 0.0017; 60 s: *p* = 0.0489), with additional differences emerging at later time points. In contrast, MIA-related differences were more apparent at later stages of the assay, particularly from 240 s onward, where fish tested in the first order remained closer to the mirror compared to those tested second (all *p* ≤ 0.005). Together, these results indicate that zebrafish behavior exhibits robust within-test temporal dynamics across assays. While the overall temporal structure of behavior is largely preserved across test sequences, test order modulates behavioral adaptation in a time– and assay-dependent manner, with effects emerging predominantly during early exploration phases but also evident at later stages in specific contexts such as aggression.

### Sex-dependent differences are restricted to the NTT

We next examined sex-dependent effects using two-way ANOVA **(Fig. 4)**. No significant main effects of sex or sex × order interactions were observed in the MIA or SP assays across all measured parameters (all *p* > 0.05), indicating that these behavioral domains are largely unaffected by sex under the present conditions. In contrast, the NTT revealed robust sex-dependent differences across multiple behavioral endpoints. Females exhibited reduced locomotor activity compared to males, as reflected by lower total distance traveled (*F* _(1,78)_ = 15.66, *p* = 0.0013) and mean speed (*F*_(1,78)_ = 15.59, *p* < 0.001). In addition, females displayed increased anxiety-like behavior, characterized by reduced time spent in the top zone (*F*_(1,78)_ = 38.64, *p* < 0.001) and decreased distance from the bottom (*F*_(1,78)_ = 45.90, *p* < 0.001).

**Fig. 4.**
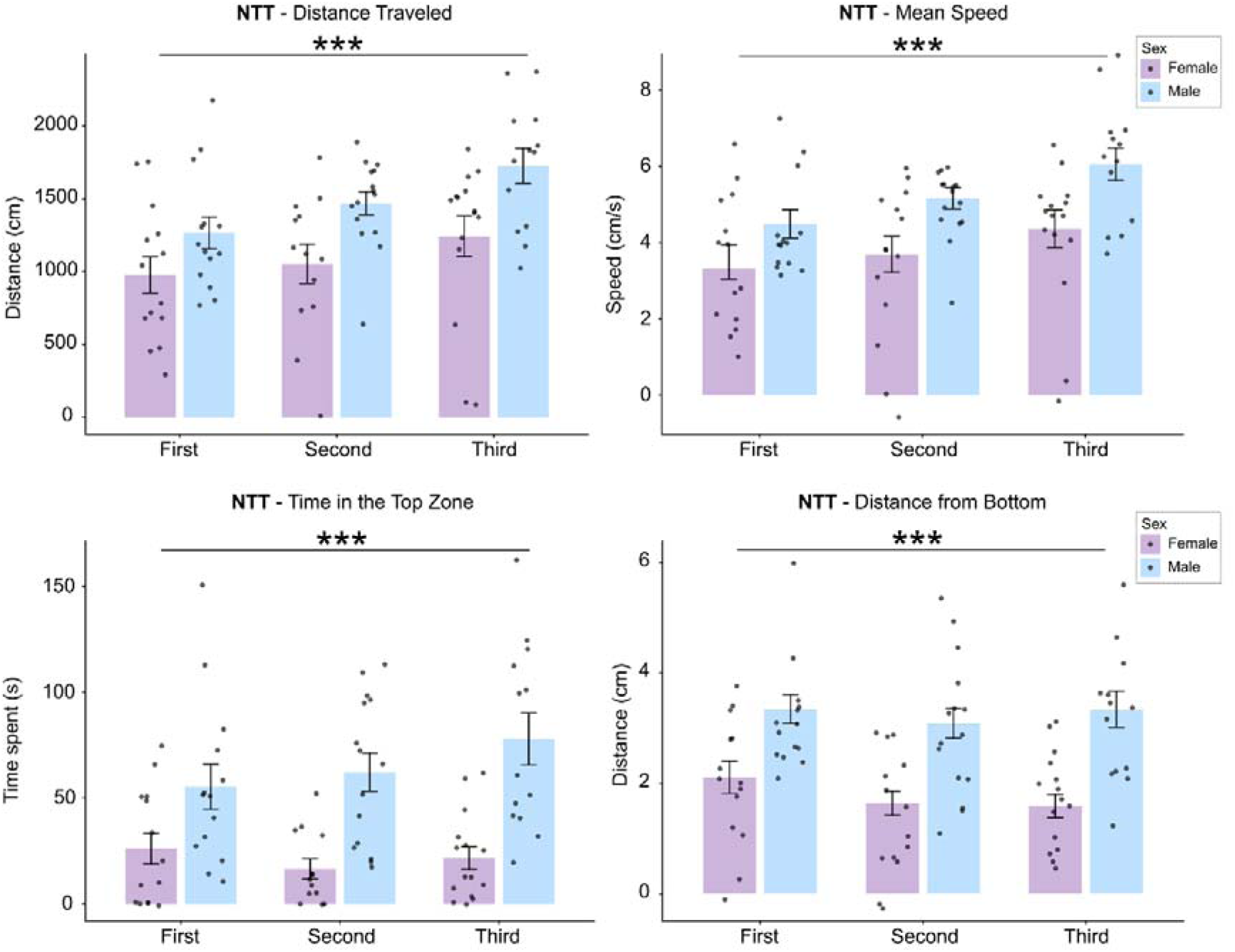
Sex-dependent effects on NTT behavioral parameters across test order. Novel tank test (NTT) behavioral parameters stratified by sex (female, male) and test position (first, second, third), including total distance traveled (cm), mean speed (cm/s), time spent in the top zone (s), and mean distance from the bottom (cm). Data are presented as mean ± S.E.M., with individual data points shown. Data were analyzed using two-way ANOVA with sex and test order as factors, followed by Tukey’s post hoc test. Asterisks indicate a significant main effect of sex (two-way ANOVA), independent of test position (*n* = 14; *p*\*\*\* < 0.001).

Significant main effects of test order within the NTT were also observed for distance traveled (*F*_(2,78)_ = 4.62, *p* = 0.013) and mean speed (*F*_(2,78)_ = 4.28, *p* = 0.017), but no significant sex × order interactions were detected (all *p* > 0.25), indicating that order effects were comparable between males and females. Together, these results demonstrate that sex-dependent differenc s are assay-specific and most pronounced in anxiety-related and locomotor behaviors measured in the NTT, whereas social and aggression-related behaviors remain largely unaffected.

### Behavioral structure is preserved across test sequences

To assess whether test sequence influenced the global structure of behavioral phenotypes, principal component analysis (PCA) was performed on standardized behavioral endpoints acro s assays **(Fig. 5)**. The first three principal components accounted for a substantial proportion of the total variance (PC1: 29.6%, PC2: 26.7%, PC3: 20.1%; cumulative: 76.4%). Inspection of variable loadings indicated that PC1 was primarily associated with social preference variables, with distance to stimulus and time near stimulus showing the largest absolute loadings (±0.545). PC2 was dominated by aggression-related measures, including distance to the mirror (–0.616) and dynamic aggression (0.588). PC3 was largely defined by NTT-derived variables, particularly distance traveled (–0.612) and distance from the bottom (–0.503), reflecting locomotor and anxiety-related dimensions. Full variable loadings for all principal components are provided in **Supplementary Table 1**.

**Fig. 5.**
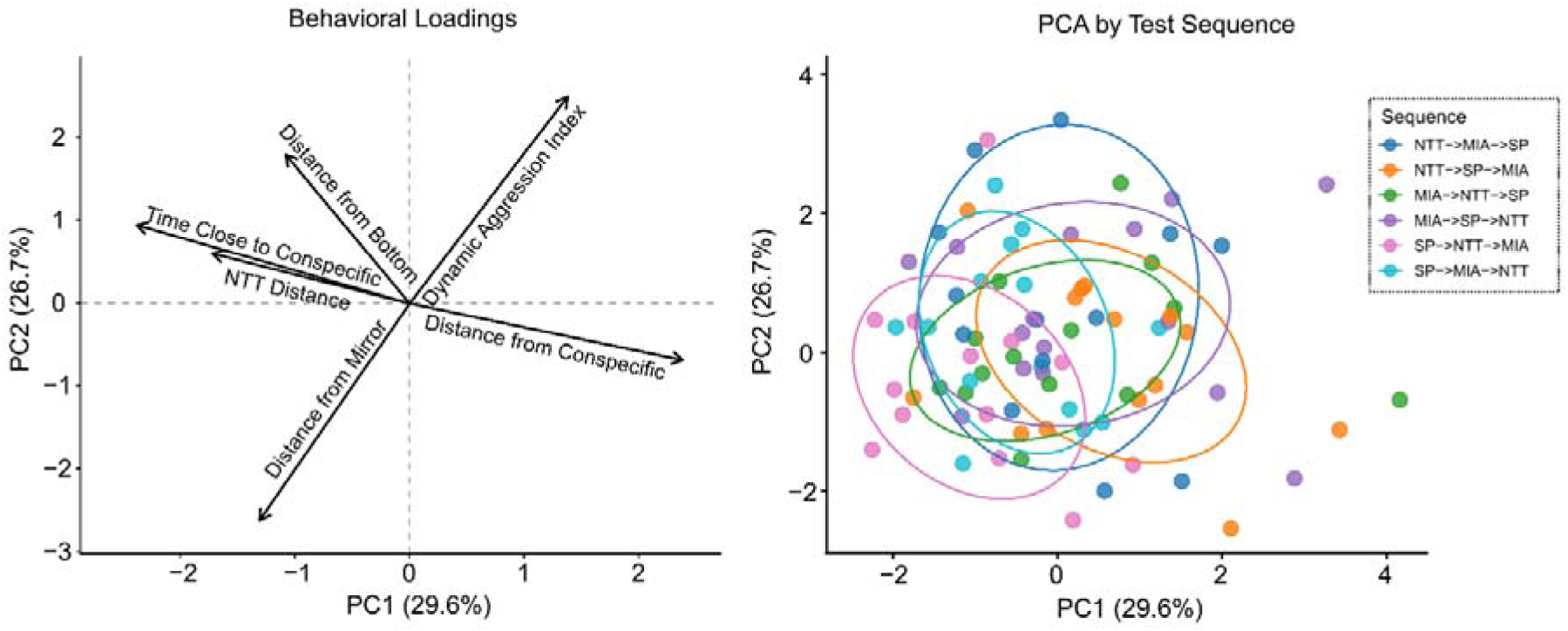
Multivariate organization of behavioral endpoints across test sequences. **(A)** Principal Component Analysis (PCA) loadings of behavioral variables across assays, showing the contribution of locomotor, anxiety-like, social, and aggression-related parameters to principal components (PC1 and PC2). **(B)** PCA score plot showing the distribution of individual animals across test sequences, with each point representing one subject and ellipses indicating group dispersion. Test sequences correspond to all permutations of the behavioral battery (NTT, MIA, SP). Data were standardized prior to PCA.

Visualization of PCA scores revealed substantial overlap across all test sequences, with no clear clustering or separation of groups in principal component space. However, multivaria e analysis detected a significant effect of test sequence on behavioral profiles (MANOVA, Pillai’s trace = 0.27, *F*_(10,156)_ = 2.45, *p* = 0.0096), indicating that sequence contributes to systematic, albeit subtle, shifts in multivariate behavioral organization. Importantly, no differences in within-group dispersion were detected (*F*_(5,78)_ = 0.58, *p* = 0.712), indicating comparable variability acro s sequences. Together, these findings indicate that, while test order does not produce clearly separable behavioral phenotypes, it exerts a modest but detectable influence on the multivariate structure of behavior across assays.

## Discussion

Sequential behavioral testing is increasingly used to characterize multidimensional phenotypes in zebrafish, yet the potential impact of test order on behavioral outcomes remains insufficiently defined. Using a fully counterbalanced three-assay battery, we demonstrate that test responses in the NTT and early responses in the SP and MIA assays. Importantly, these localized effects did not result in major disruption of the overall multivariate behavioral structure, supporting the use of multidomain behavioral batteries under baseline conditions. These findings extend previous observations suggesting that sequential behavioral testing can be implemented without substantial interference in naive zebrafish^22^ while emphasizing the importance of considering temporal and assay-specific carryover effects in experimental design.

The most robust order-dependent effects were observed in the NTT, where animals tested later exhibited increased locomotion. This pattern is consistent with carryover effects driven by prior exposure, a factor increasingly recognized in behavioral battery designs^18,20,21^. Two non-exclusive mechanisms may underlie this effect. First, prior exposure to experimental contexts may reduce novelty-induced suppression, facilitating increased exploratory drive. This interpretation is consistent with the ethological basis of the NTT, in which zebrafish initially display bottom-dwelling behavior followed by progressive exploration during habituation^1,3,23,24^. Second, prior testing^25^ and repeated handling^26^ may elevate arousal or stress responsiveness, leading to generalized increases in locomotor output. Given that locomotor parameters are particularly sensitive to stress and environmental conditions, including handling-related factors^25,27^, these findings suggest that locomotion is especially susceptible to cumulative experimental exposure. Notably, comparable effects were not observed across the other assays, indicating that the NTT may represent a particularly sensitive paradigm for detecting subtle carryover effects.

Temporal analysis further demonstrated that these effects were dynamic and phase-dependent rather than static. In the NTT, animals exposed to prior tests exhibited reduced distance from the bottom during the initial phase of the assay, consistent with enhanced anxiety-like behavior at test onset. However, these differences rapidly diminished over time as animals habituated to the environment, eventually displaying increased overall locomotion relative to animals tested first. These findings further support the interpretation that prior assay exposure modifies adaptive responses to novelty, producing enhanced early anxiety-like behavior followed by accelerated exploratory engagement during habituation. Because these early-phase effects were not consistently detected using conventional aggregate 5-min endpoints, subtle influences of sequential behavioral testing may remain undetected when relying exclusively on standard summary measures^28^.

In contrast, animals tested second or third in the MIA assay exhibited reduced proximity to the mirror over time, suggesting decreased aggressive-like engagement. Similarly, fish tested later in the SP assay showed reduced early social approach, remaining farther from conspecifics during the initial phase of testing. These findings suggest that sequential testing induces context-dependent carryover effects on motivational and social processes, potentially reflecting cumulative handling or altered responsiveness to social stimuli. Cumulative stress exposure may also contribute to these behavioral dynamics, as stress-related states are known to modulate social and aggression-related behaviors in zebrafish^15,29,30^, although physiological stress markers were not directly assessed in the present study. Notably, these effects were most evident during early time bins and may therefore be underestimated using conventional aggregate endpoints. This consideration may be particularly relevant for SP paradigms, which are frequently analyzed using short recording windows or first-minute interactions^31–34^, where transient differences in early social engagement could substantially influence final behavioral outcomes. Previous methodological discussions have also highlighted substantial variability in habituation and interaction durations across zebrafish SP protocols, emphasizing the need for greater temporal standardization and characterization of behavioral dynamics during testing^4^. Importantly, despite these early-phase effects, overall behavioral measures across the full 5-min sessions remained largely preserved, indicating that carryover effects in these assays are modest and primarily time-dependent. Together, our findings support the incorporation of time-resolved analyses, rather than reliance on restricted early recording windows alone, to better capture dynamic behavioral adaptations during behavioral testing.

Sex differences were also investigated, and these effects were assay-specific, being primarily observed in the NTT, where females displayed increased anxiety-like behavior compared to males. This finding is consistent with previous data showing that female zebrafish exhibit enhanced anxiety-like responses across assays such as the NTT and LDT, characterized by increased bottom-dwelling and dark preference^35^. These observations have been further supported by large-scale multi-laboratory data demonstrating a consistent tendency toward higher anxiety-like behavior in females^27^. However, no sex effects were detected in the MIA or SP assays, suggesting that social and aggression-related behaviors were comparatively less sensitive to baseline sex differences under the present conditions. These results highlight the importance of considering sex as a biological variable in zebrafish behavioral studies while also indicating that sex-dependent effects may be strongly domain-specific.

We further assessed whether test order influences multidimensional behavioral organization. At the multivariate level, behavioral variation was structured along distinct functional axes corresponding to locomotor, anxiety-related, and social domains. This integrative approach is increasingly recognized as essential for capturing complex behavioral phenotypes and improving reproducibility across studies^36–38^. Despite detectable univariate effects of test order, there was no clear segregation of behavioral profiles across test sequences, indicating that sequential testing does not induce major shifts in global behavioral organization. Instead, test order contributed to subtle modulations within an otherwise stable phenotypic space, suggesting that behavioral batteries remain a robust approach for multidimensional phenotyping. This is particularly relevant in the context of ongoing reproducibility challenges in zebrafish behavioral neuroscience^39^, where methodological variability, including differences in testing protocols and experimental sequence, has been identified as a major source of inconsistency^19^.

Although we clearly demonstrate that test battery used here elicits modest effects on zebrafish behavior, several limitations should be considered. Although a fully counterbalanced design was employed, the present study does not establish an optimal or standardized test sequence, and different assay combinations or intervals between tests may yield distinct outcomes. Notably, the NTT emerged as the most sensitive assay to test order, exhibiting detectable carryover effects on both locomotor and early anxiety-like responses, whereas effects in MIA and SP were largely restricted to transient, time-dependent changes that did not impact overall behavioral endpoints. These findings suggest that carryover effects are assay-dependent, primarily influencing early behavioral dynamics, and may therefore be underestimated when relying solely on conventional summary measures. In addition, repeated handling and test exposure may contribute to cumulative stress, which was not directly assessed in the present study. Future work incorporating physiological measures, such as cortisol levels or other stress-related biomarkers, will be important to better characterize the mechanisms underlying these carryover effects. Furthermore, studies including additional behavioral paradigms, longer inter-test intervals, or repeated testing designs will help define the boundaries and generalizability of these effects. In addition, the present study was performed using short-fin wild-type zebrafish, and potential differences related to zebrafish populations or behavioral phenotypes should be considered in future behavioral battery studies. Studies should also evaluate how test order interacts with pharmacological manipulations, as prior evidence suggests that drug effects may vary depending on test sequence^22^, highlighting an additional layer of experimental complexity.

Here, our findings have important implications for experimental design. While sequential behavioral testing can be implemented without major confounds under baseline conditions, test order may still influence specific behavioral endpoints and temporal response dynamics. Therefore, careful standardization and reporting of test sequence remain important for ensuring reproducibility and cross-study comparability^19^. Importantly, the preservation of the overall multivariate behavioral structure across test sequences supports the use of multidomain behavioral batteries for comprehensive phenotypic characterization. By enabling multiple behavioral domains to be assessed within the same individuals, this approach aligns with the principles of Replacement, Reduction, and Refinement (3Rs), maximizing the information obtained from each experimental cohort while reducing animal use.

## Conclusions

In conclusion, test order exerts modest, transient, and domain-specific effects on zebrafish behavior, primarily influencing locomotor and early anxiety-like responses in the NTT, as well as early social engagement in the SP assay, without substantially altering global behavioral organization. Several order-dependent effects were most evident during early phases of testing and were not consistently detected using conventional aggregate 5-min endpoints, indicating that time-resolved analyses provide increased sensitivity for identifying subtle carryover effects during sequential behavioral testing. Overall, these findings support the use of multidomain behavioral batteries in zebrafish while emphasizing the importance of temporal analyses and standardized reporting of test sequence to improve reproducibility and cross-study comparability, particularly for novelty-based paradigms such as the NTT.

## Material and Methods

### Animal husbandry

Adult wild-type zebrafish (*n* = 84; 4–6 months old; ∼50:50 female:male ratio) were obtained from a local supplier (Hobby Aquários, RS, Brazil). Sample size was determined a priori assuming a medium effect size (f = 0.25), α = 0.05, and power = 0.75 for a six-group ANOVA design evaluating behavioral differences across test orders, resulting in 14 animals per group. The calculation was based on total distance traveled in the novel tank test (NTT), which was designated as the primary outcome measure. Fish of the short-fin phenotype were maintained under constant filtration and aeration in 40-L tanks filled with non-chlorinated water (2 fish/L). Water temperature was maintained at 27 ± 1 °C, pH 7.0–7.2, dissolved oxygen at 6.0 ± 0.1 mg/L, and total ammonia < 0.01 mg/L. Animals were acclimated to laboratory conditions for at least 14 days prior to experimentation and fed three times daily with commercial granulated food (Daily Gran, OceanTech, Brazil). Fluorescent lamps provided a 14:10 h light/dark photoperiod (lights on at 07:00). Behavioral testing was conducted between 09:00 and 16:00, and tank water was renewed between subjects^40^.

### Experimental design

To evaluate the effects of behavioral battery order, adult zebrafish were sequentially exposed to three behavioral assays: the NTT, SP and MIA, with immediate transfer between assays using a soft mesh net. Animals were randomly selected from holding tanks based on visual sex identification and randomly assigned to one of six experimental groups corresponding to all possible test orders **(Fig. 1)**. Each fish was tested in all three assays within a single experimental session, with the order determined by group allocation. This fully counterbalanced design allowed each assay to occur in first, second, or third position across groups. Experiments were performed across multiple days (4 experimental batches) using randomized testing order to minimize batch effects. Behavioral sessions were video recorded for offline analysis. The NTT was recorded using a frontal camera positioned in front of the apparatus to capture vertical exploration. The SP and MIA tests were recorded from a top-view perspective. All videos were recorded using a Logitech C920 Pro HD camera (1920 × 1080 resolution, 30 frames/s). Videos were exported in MP4 format and analyzed offline using idtracker.ai for automated tracking of individual fish^41^. Tracking outputs with frame positional coordinates (X–Y) were used to extract to construct behavioral endpoints for each assay. All analysis scripts and processed datasets used for behavioral quantification are publicly available at the project GitHub repository (github.com/BarbaraDFontana/Zebrafish_Behavioral_Battery_Analysis). During behavioral quantification, animals were identified only by numerical IDs, and group allocation information was stored separately until statistical analyses were performed. Camera position, distance, and lighting conditions were kept constant across all experimental sessions to ensure consistency in tracking performance. No animals were excluded from the analysis. Following completion of the behavioral battery, animals were anesthetized using rapid cooling (2–4 °C) and euthanized by decapitation. Sex was initially determined based on external morphological characteristics (body shape and coloration) and subsequently confirmed post-mortem by gonadal inspection^42^. Identification was performed by two trained observers.

### Novel Tank Diving Test (NTT)

The NTT assessed anxiety-like behavior by quantifying vertical distribution, as zebrafish typically bottom-dwell in novel environments and progressively explore upper regions as anxiety decreases^1,43^. Fish were individually placed in the experimental tank (25 cm length × 15 cm height × 5 cm width; 10 cm water column depth) and recorded for 5 min. Behavioral endpoints extracted from positional tracking included total distance traveled (cm), mean speed (cm/s), immobility (s), time spent in the top zone (s), and distance from bottom (cm). Immobility was defined as periods of low velocity sustained for ≥1 s, using a velocity threshold (< 1 cm/s) applied to frame-by-frame speed estimates^23^. The same analysis of immobility was used across behavioral tests.

### Mirror-induced aggression (MIA)

The MIA test assesses aggression-related behavior by using the zebrafish tendency to display agonistic interactions toward their mirror reflections. The test was performed as previously described^8,44^. Fish were individually placed in a rectangular tank (25 cm length × 15 cm height × 10 cm width; 10 cm water column depth) containing an inclined mirror (22.5°) positioned on one side to elicit aggressive responses toward the reflected image. All other tank sides were covered with opaque partitions to minimize external visual interference. Animals were recorded for 5 min, and locomotor parameters were extracted from positional tracking data, including total distance traveled (cm), mean speed (cm/s), and immobility (s). Aggression-related behavior was quantified using continuous spatial metrics relative to the mirror plane. The perpendicular distance to the mirror was computed at each frame, with shorter distances interpreted as increased aggressive motivation. To account for the inclined geometry of the mirror and variation in fish position along the vertical axis, a tilt-corrected effective distance to the mirror was calculated by incorporating normalized vertical position. A dynamic aggression index was then defined as the inverse of this effective distance and normalized within each trial, yielding a continuous measure of proximity-based aggressive engagement that increases with sustained close positioning to the mirror. Behavioral endpoints included the mean distance to the mirror and the mean dynamic aggression index.

### Social Preference Test (SP)

The SP test assesses social behavior by quantifying the preference of an individual fish for proximity to a conspecific stimulus. The test was performed based on previous protocols^4,15,31^. Fish were individually placed in an experimental tank (25 cm length × 15 cm height × 10 cm width; 10 cm water depth) adjacent to a stimulus tank containing six naïve conspecifics and a blue area (opposite side). Behavioral testing lasted 5 min, with fish transferred directly from housing tanks or preceding assays within the behavioral battery. Conspecific stimulus fish were drawn from a separate housing tank and were not subjected to any behavioral testing. Behavior was quantified using positional tracking and spatial metrics relative to the stimulus location. Locomotor parameters included total distance traveled (cm), mean swimming speed (cm/s), and immobility (s). Social preference was assessed as the distance to conspecifics (cm), computed at each frame, with lower values indicating increased social approach. In addition, the tank was subdivided into four equal virtual zones, each representing 25% of the total tank length, and the time spent in the zone close to the conspecific stimulus was used as a complementary measure of social preference. Animals spent minor time in the blue area; therefore, this zone was not included in the data analysis.

### Statistical analysis

Behavioral data were analyzed using R statistical computing environment (version 4.4.0). Analyses were performed on endpoints derived from 5-min recordings, unless otherwise specified. Model assumptions were assessed prior to statistical analyses. Model assumptions were assessed prior to analysis, including evaluation of residual normality and homogeneity of variances. Data met the assumptions required for parametric analyses. To assess the effect of test order on behavioral endpoints, one-way analyses of variance (ANOVA) were conducted separately for each assay (NTT, MIA, and SP), with test position (first, second, or third) as the independent factor. Two-way ANOVA models were additionally applied to evaluate the effects of test order and prior exposure (“before” condition), as well as their interaction. Sex-dependent effects were also examined using two-way ANOVA with sex and test order as fixed factors. Significant effects were followed by Tukey’s honestly significant difference (HSD) post hoc tests with adjustment for multiple comparisons. To evaluate behavioral dynamics across time, data were segmented into 30-s bins over the 5-min recording period and analyzed using linear mixed-effects models, with test order and time as fixed factors and subject (ID) included as a random intercept to account for repeated measures. Statistical significance was assessed using Type III ANOVA with Satterthwaite’s approximation for degrees of freedom. Post hoc comparisons across test order within time bins were performed using estimated marginal means. To assess whether test sequence influenced the global structure of behavioral phenotypes, a Principal Component Analysis (PCA) was performed on standardized behavioral endpoints across assays. Group differences in multivariate behavioral space were subsequently evaluated using Multivariate Analysis of Variance (MANOVA), with test sequence as the independent factor and PCA scores as dependent variables. Data are presented as mean ± standard error of the mean (SEM), with individual data points shown where appropriate. Statistical significance was set at *p* < 0.05.

## Ethics Statement

All procedures were conducted in accordance with the ARRIVE Guidelines 2.0 (see **Supplementary Material**). Experiments were approved by the Institutional Animal Use Ethics Committee (CEUA/UFSM, protocol number 412110722), in compliance with National Institute of Health Guide for Care and Use of Laboratory Animals guidelines for the care and use of experimental animals.

## Data availability

The dataset and analysis scripts used in this study are publicly available at: github.com/BarbaraDFontana/Zebrafish_Behavioral_Battery_Analysis.

