## Supplementary material for "Behavioral test batteries induce transient, domain-specific effects while preserving global phenotypic structure in zebrafish": Table S1.

**Supplementary Table 1.** Variable loadings for each principal component.

| **Variable** | **PC1** | **PC2** | **PC3** | **PC4** | **PC5** | **PC6** |
| --- | --- | --- | --- | --- | --- | --- |
| Distance from bottom (NTT) | -0.247 | 0.423 | -0.503 | -0.628 | -0.214 | 0.259 |
| Distance traveled (NTT) | -0.394 | 0.142 | -0.612 | 0.522 | 0.189 | -0.377 |
| Distance to mirror (MIA) | -0.300 | -0.616 | -0.223 | 0.205 | -0.173 | 0.639 |
| Dynamic aggression (MIA) | 0.315 | 0.588 | 0.016 | 0.519 | -0.137 | 0.517 |
| Distance to stimulus (SP) | 0.545 | -0.161 | -0.416 | -0.142 | 0.671 | 0.183 |
| Time near stimulus (SP) | -0.545 | 0.223 | 0.387 | -0.048 | 0.648 | 0.285 |
